# Multiple optimal phenotypes overcome redox and glycolytic intermediate metabolite imbalances in *Escherichia coli pgi* knockout evolutions

**DOI:** 10.1101/342709

**Authors:** Douglas McCloskey, Sibei Xu, Troy E. Sandberg, Elizabeth Brunk, Ying Hefner, Richard Szubin, Adam M. Feist, Bernhard O. Palsson

## Abstract

A mechanistic understanding of how new phenotypes develop to overcome the loss of a gene product provides valuable insight on both the metabolic and regulatory function of the lost gene. The *pgi* gene, whose product catalyzes the second step in glycolysis, was deleted in a growth optimized *Escherichia coli* K-12 MG1655 strain. The knock-out (KO) strain exhibited an 80% drop in growth rate, that was largely recovered in eight replicate, but phenotypically distinct, cultures after undergoing adaptive laboratory evolution (ALE). Multi omic data sets showed that the loss of *pgi* substantially shifted pathway usage leading to a redox and sugar phosphate stress response. These stress responses were overcome by unique combinations of innovative mutations selected for by ALE. Thus, we show the coordinated mechanisms from genome to metabolome that lead to multiple optimal phenotypes after loss of a major gene product.

**Importance:** A mechanistic understanding of how new phenotypes develop to overcome the loss of a gene product provides valuable insight on both the metabolic and regulatory function of the lost gene. The *pgi* gene, whose product catalyzes the second step in glycolysis, was deleted in a growth optimized *Escherichia coli* K-12 MG1655 strain. Eight replicate adaptive laboratory evolution (ALE) resulted in eight phenotypically distinct endpoints that were able to overcome the gene loss. Utilizing multi-omics analysis, we show the coordinated mechanisms from genome to metabolome that lead to multiple optimal phenotypes after loss of a major gene product.

## Introduction

The flux split between upper glycolysis and the oxidative pentose phosphate pathway (oxPPP) at the glucose 6-phosphate (G6P) node is a major determinant of the flux state of a cell’s core metabolic network. Loss of Phosphoglucose Isomerase (PGI), encoded by *pgi*, induces detrimental physiological consequences (1–5). Removal of *pgi* generates an imbalance in glycolytic intermediates from the loss of upper glycolytic flux that leads to a loss of fitness, and induces a sugar phosphate stress response. The sugar phosphate stress response involves the actions of both small RNAs (sRNAs) and transcription factors (TFs) that induce transcription level changes aimed at alleviating the imbalance (6–8). Removal of *pgi* also generates an overabundance of NADPH and redox imbalance by redirecting glycolytic flux into the oxPPP. NADPH provides reducing equivalents for biosynthesis of lipids, cholesterol, and other macromolecules. In addition, NADPH plays an important role in reactive oxygen species (ROS) detoxification by regenerating reduced glutathione (gthrd) (9). Increased availability of NADPH in *pgi-* backgrounds has proven useful in various biotechnology applications in order to increase cofactor and heterologous pathway production (1, 2, 10).

Adaptive laboratory evolution (ALE) of *pgi* mutants have been carried out to better understand the physiological changes required to overcome genetic perturbation (3, 4). ALE is an experimental method that introduces a selection pressure (e.g., growth rate selection) in a controlled environmental setting (11–13). Using ALE, organisms can be perturbed from their evolutionary optimized homeostatic states, and their re-adjustments can be studied during the course of adaptation to reveal novel and non intuitive component functions and interactions (14). Previous ALEs of *pgi* mutants have demonstrated a re-wiring of central metabolic fluxes (4) and diversity in endpoint physiological phenotypes (3). The diversity in endpoint physiological phenotypes is directly attributed to the existence of alternate optimal metabolic and regulatory network states that can achieve the same physiological function (3). However, the mechanisms and coordination of the regulatory and metabolic network required to produce physiologically distinct, yet fit, phenotypes is not well understood. In addition, these studies were conducted with a starting strain that was not previously optimized to the growth conditions of the experiment. This confounds the interpretation of the experimental results because adaptations to the growth conditions and loss of the gene occur simultaneously.

The consequence of the loss of a major metabolic gene can be studied at the systems level through the integration of multi-omics data sets (i.e., metabolomics, fluxomics, proteomics, and transcriptomics) to gain deeper insight into the function of the gene in the context of the biological system as a whole. Previous work has found that the metabolic network is robust to perturbations through adjustments made at the regulatory level that coordinate re-routing of flux with enzyme level (5, 15). While these studies reveal insights to the immediate response of gene loss, the adaptive changes required to overcome the loss were not explored. In addition, improvements in -omics data acquisition and analysis methods could improve and reveal new relationships between changes in -omics data at one layer of the system to another.

In this study, a combination of experimental design (i.e., starting with a strain that was pre evolved on glucose M9 minimal media) and systems analysis from multi-omics data was used to mechanistically investigate how multiple phenotypes can overcome the loss of *pgi*. First, the reduction in fitness after the *pgi* KO was found to be attributed to malfunctions in the regulatory and metabolic network that were incapable of handling the redox and glycolytic intermediate metabolite imbalance induced from major shifts in central metabolic flux. Second, all evolved *pgi* KO lineages regained a substantial portion of fitness, but displayed unique physiologies. Third, the recovery in fitness was made possible by mutations that were selected for by ALE that altered the transcription regulatory network (TRN) and metabolic fluxes to alleviate the redox and sugar phosphate imbalance. These regulatory and metabolic alterations were unique across all endpoints, which lead to the emergence multiple optimal phenotypes.

## Results

### Diversity in ALE endpoint phenotypes points to multiple optimal selection outcomes

To eliminate the confounding variable of adaptation to the growth conditions of the experiment, a wild-type *E. coli* K-12 MG1655 strain previously evolved under glucose minimal media at 37°C(16) (denoted as “Ref”) was selected as the starting strain (Fig. 1A). This selection was made to separate changes caused by adaption to the loss of a gene product from those caused by adaption to the growth conditions of the experiment.

**Fig. 1.**
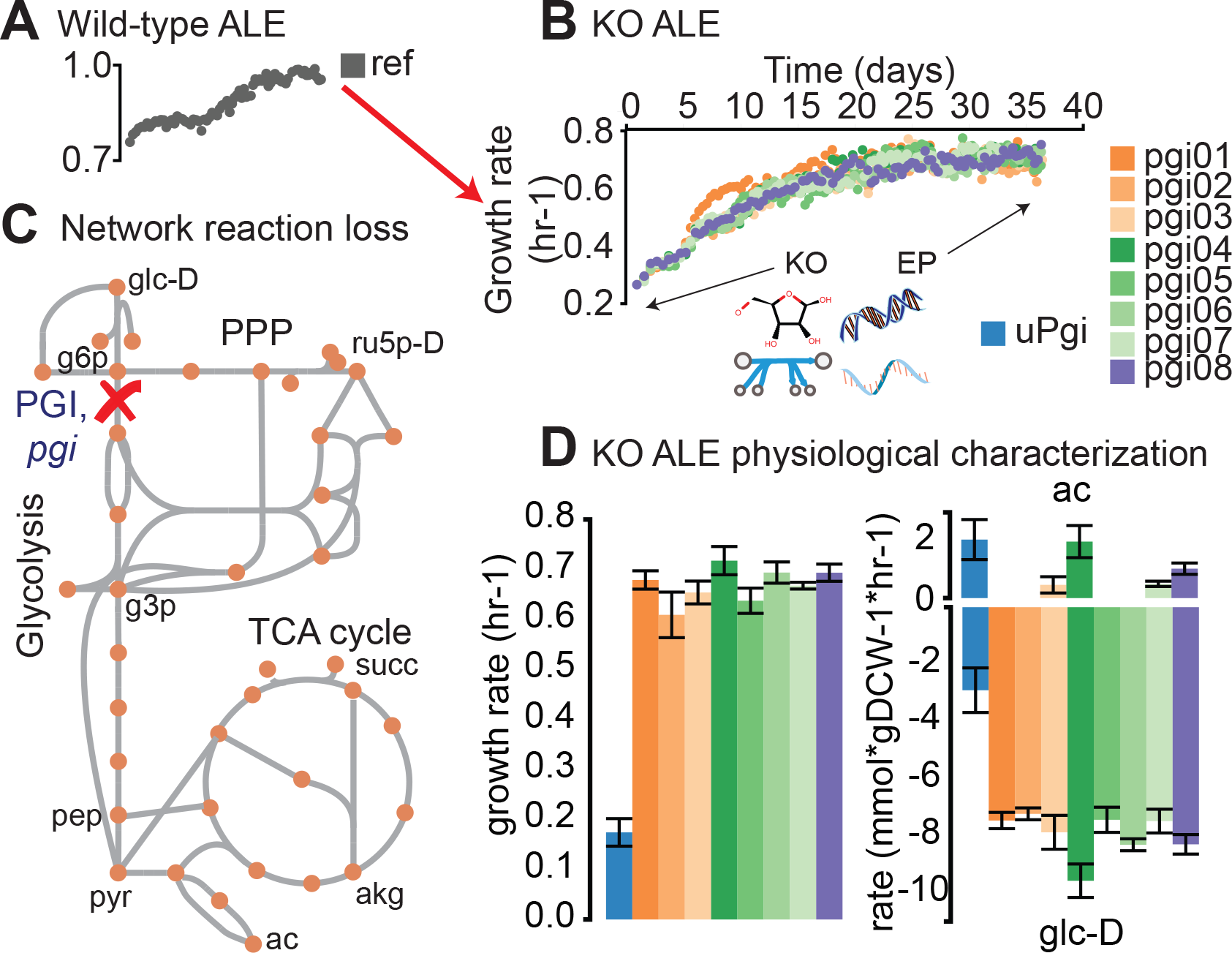
Evolution of knockout (KO) strains from a pre-evolved (i.e., optimized) wild-type strain. A) Wild-type (wt) *E. coli* (MG1655 K-12) was previously evolved on glucose minimal media at 37°C(16). An isolate from the endpoint of the evolutionary experiment was selected as the starting strain for subsequent KO of *pgi* and adaptive laboratory evolution (ALE). B) Adaptive laboratory evolution trajectories of the evolved knockout lineages. -Omics data collected from the fresh KO, and end-point lineages included metabolomics, fluxomics, physiology, DNA resequencing, and transcriptomics. C) Phosphoglucose isomerase (PGI) was disabled by the gene KO. PGI is the first step in glycolysis and converts glucose 6 phosphate (g6p) to fructose 6 phosphate (f6p). D) Growth rate and glucose (glc-D) uptake and acetate (ac) excretion rates for unevolved KO (uPgi) and evolved KOs (ePgi). Error bars denote 95% confidence intervals from biological triplicates.

PGI (*pgi*, phosphoglucose isomerase) was removed from Ref to generate strain uPgi (denoted “unevolved *pgi* knockout strain”) (Fig. 1B). The loss of *pgi* resulted in an 81% loss in growth rate (Fig. 1 C, D). Eight uPgi independently inoculated starting cultures were simultaneously evolved on glucose minimal media at 37°C in an automated ALE platform (16, 17) denoted “evolved *pgi* knockout strains” or “ePgi” (Fig. 1C). A statistically significant increase in final growth rate (Student’s t-test, pvalue<0.05) was found in all ALE endpoints of the ePgi lineages (ave±stdev 284±20% increase in growth rate) compared to uPgi (Fig. 1D). Metabolomics, fluxomics, transcriptomics, genomics, and phenomics data was collected from exponentially growing cultures inoculated in triplicate from Ref, uPgi, and each of the 8 independently evolved end-point lineages (ePgi01-08).

Statistically significant variability in growth rate, acetate secretion, and glucose uptake rate were found in ePgi strains (Fig. 1D, Table S2). Specifically, replicates 3, 4, 7, and 8 excreted acetate.

Replicate 4 in particular had acetate secretion levels similar to uPgi, and the highest growth and glucose consumption rate of all endpoints. It should be emphasized that all

These results raised two defining questions: What metabolic and regulatory changes occurred to allow for a large regain in fitness without the use of upper glycolysis? How were a diversity of end point physiologies capable of overcoming the loss of PGI? To answer these questions, Intracellular metabolite levels, gene expression levels, flux levels, and genomic mutations were measured for the Ref, uPgi, and ePgi strains (Tables S3-8).

### PGI KO shifted metabolic flux

Genome-scale MFA(18) found significant shifts in flux splits throughout central metabolism in response to the loss of PGI (Fig. 2, Table S6). Note that all fluxes discussed in the main text passed observability criteria as described previously (18). Flux splits included the distribution of flux through the oxidative Pentose Phosphate Pathway, oxPPP, (EDD, GND, and PGL), flux through the non-oxidative branch (nonOxPPP), flux around the anaplaerotic reactions (PPC, PPCK, MALS, ME1, and ME2), flux into the TCA cycle or towards acetate secretion (CS, ACt2rpp, and PDH), and flux through the lower glyoxylate shunt or through the lower TCA cycle (ACONTb, ICDHyr, and ICL). A massive 12601.3% increase in flux per mol of glucose through the ED Pathway (EDD) was found, while a minor 15.6% drop in flux per mole of glucose through GND was found in uPgi compared to Ref. Redistribution of flux through the nonOxPPP was found. A 79% increase in flux per mole of glucose into the TCA through CS was offset by an 87% drop in flux per mole of glucose into the TCA through PPC. A minor 6% drop in acetate secretion per mole of glucose was found. A 380.8% increase in flux per mole of glucose through the glyoxylate shunt was found.

**Fig. 2.**
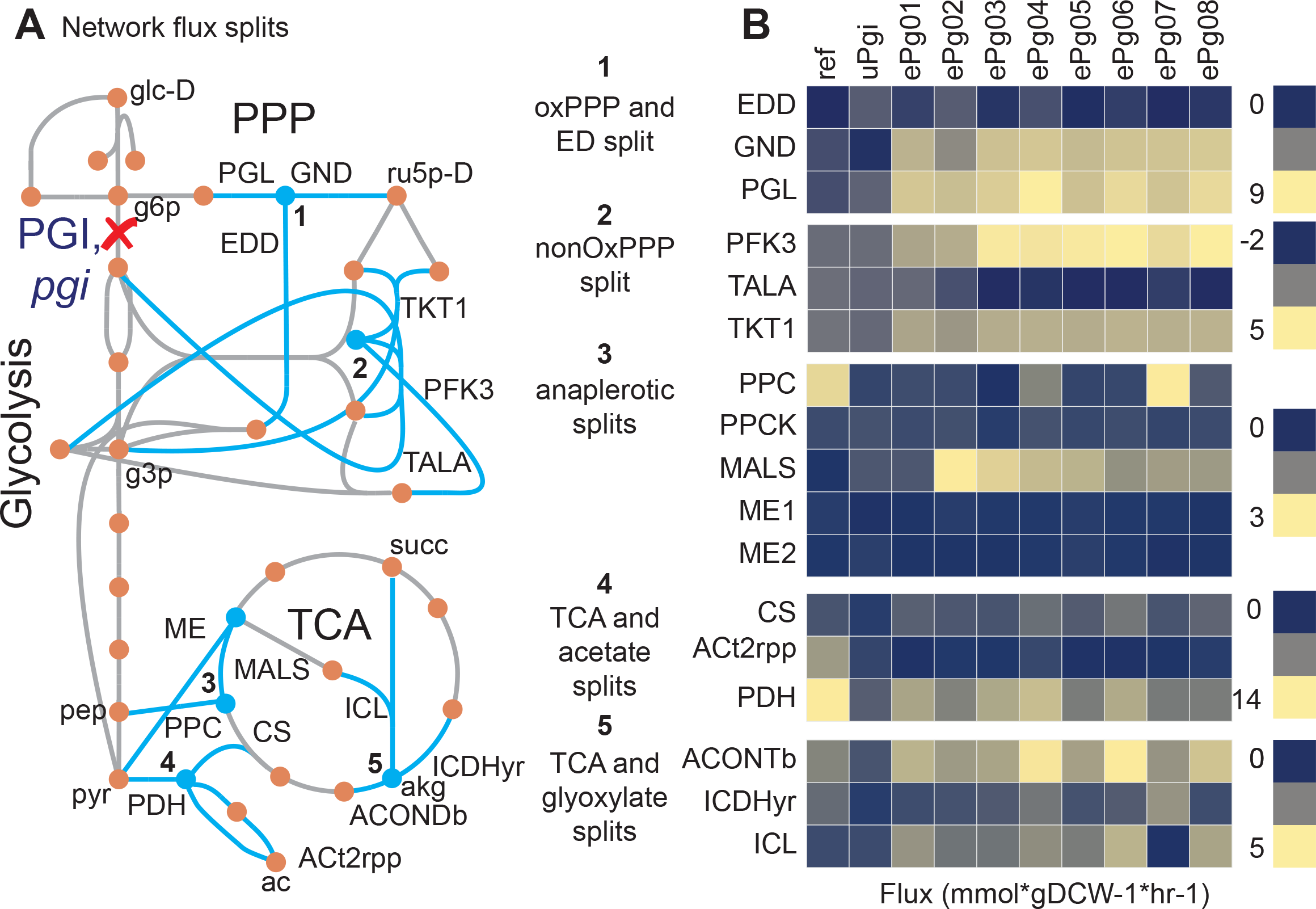
Changes in flux splits pre- and post- adaptive evolution. A) Network diagram with reactions involved in flux splits annotated. Reactions included phosphogluconate dehydratase (EDD), 6-phosphogluconate dehydrogenase (GND), 6-phosphogluconolactonase (PGL), phosphoenol pyruvate carboxylase (PPC), phosphoenolpyruvate carboxylase kinase (PPCK), malate dehydrogenase (MALS), NADP-dependent malic enzyme (ME1), NAD-dependent malic enzyme (ME2), citrate synthase (CS), accetate secretion (ACt2rpp), pyruvate dehydrogenase (PDH), aconitase (ACONTb), isocitrate dehydrogenase (ICDHyr), and isocitrate lyase (ICL). B) Measured absolute fluxes for Ref, uPgi, and ePgi strains. Values are derived from averages taken from triplicate cultures that were analyzed in duplicate (n=6).

### Perturbed glycolytic intermediates generated a sugar phosphate stress response

Shifts in central metabolic fluxes imbalanced central metabolic intermediate metabolite levels. LC-MS/MS was used to quantify the absolute metabolite concentrations of glycolytic intermediates, PPP, and TCA cycle intermediates (Table S3); transcriptomics was used to quantify the relative shifts in genes targeted by transcription factors (Table S4-5). All measured glycolytic and PPP intermediates changed significantly in uPgi compared to Ref (Fig. 3, Table S3). In particular, an approximate fivefold change in glucose 6-phosphate (g6p) was found in uPgi compared to Ref (Fig. 4). Abnormal elevations in g6p and an imbalance of the glycolytic intermediates in uPgi were found to induce the sugar phosphate toxicity response transcription factor (TF) SgrR(6–8). SgrR is thought to bind hexose phosphates and induce the expression of the small RNA *sgrS (6–8)*(Fig. 4D-E). *sgrS* was highly overexpressed in uPgi compared to Ref and ePgi strains. *SgrS* transcriptionally regulates a number of genes including the *pur* regulon, *ptsG*, and genes involved in biofilm formation and curli formation (7, 8, 19–21).

**Fig. 3.**
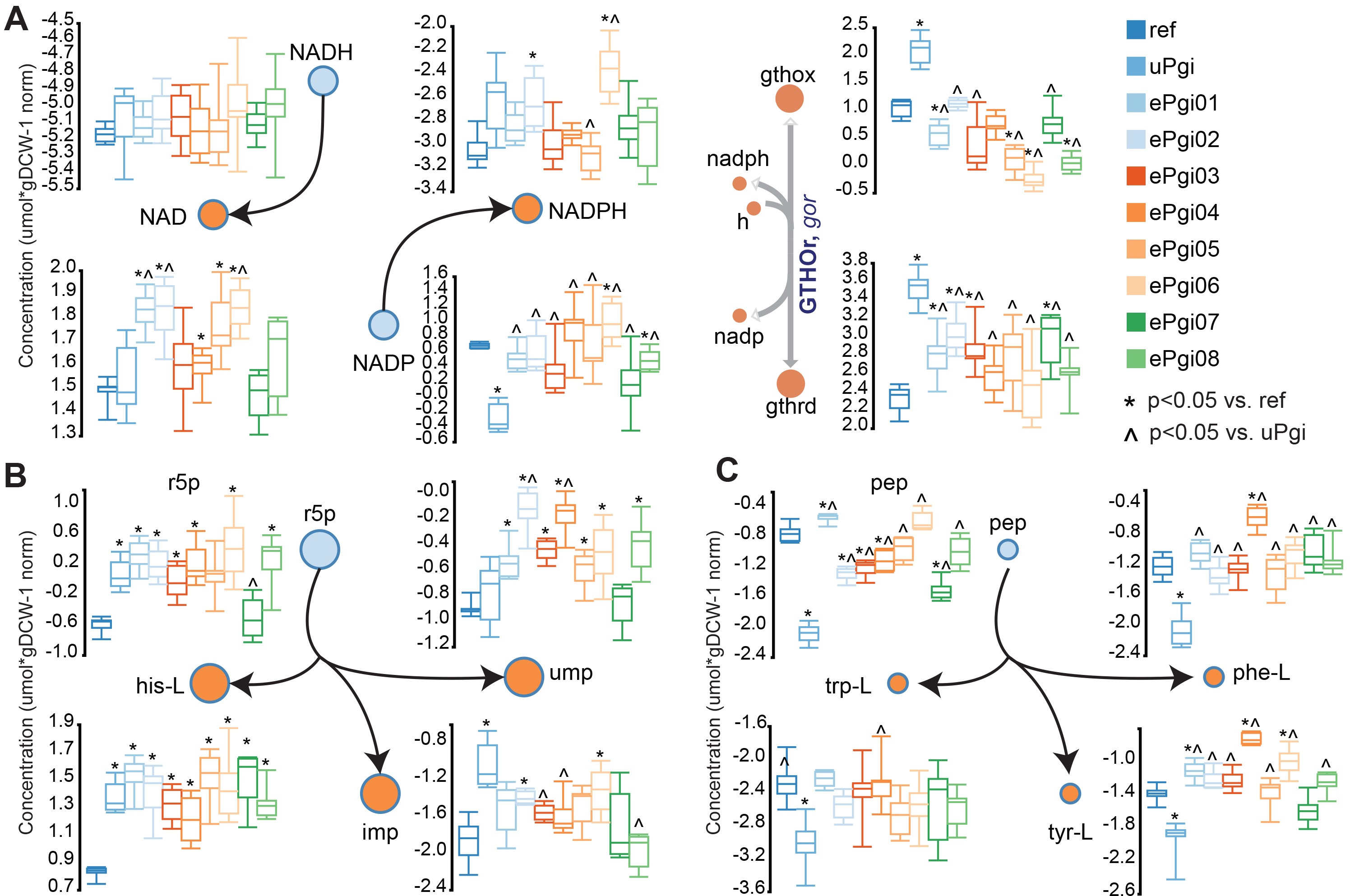
An imbalance in redox carriers. A) Box and whiskers plots of log-normalized absolute metabolite levels (umol*gDCW-1 norm) of the redox carriers NAD(P)(H) and the reduced (gthrd) and oxidized (gthox) Glutathiones. Network diagram of the interconversion of nadh to nad, nadp to nadph, and gthox and nadph to gthrd and nadp. An imbalance in glycolytic and PPP intermediates, and their downstream biosynthetic components. A) Schematic of the connection between the PPP precursor ribose 5 phosphate (r5p) and downstream amino acid and nucleotides L-histidine (his-L), Inosine Monophosphate (IMP), and Uridine Monophosphate (UMP). Box and whiskers plots of absolute metabolite levels of r5p, his-L, imp, and ump. B) Schematic of the connection between the glycolytic precursor Phosphoenol Pyruvate and downstream aromatic amino acids L-Tryptophan (trp-L), L-Tyrosine (tyr-L), and L-Phenylalanine (phe-L). Box and whiskers plots of pep, trp-L, tyr-L, and phe-L. Values are derived from averages taken from triplicate cultures that were analyzed in duplicate (n=6).

**Fig. 4.**
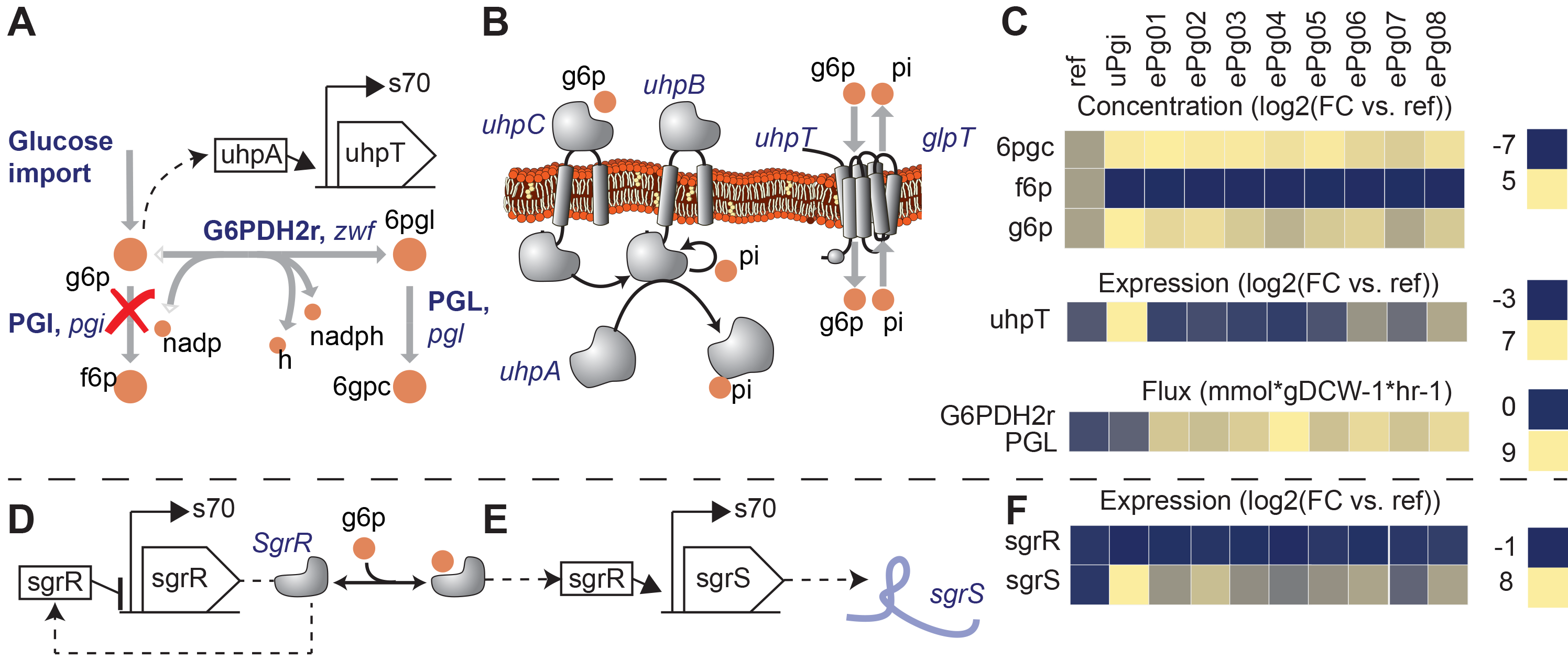
KO of PGI led to a hexose phosphate toxicity response. The magnitude of g6p in the initial knockout led to a deleterious cycle whereby leakage of hexose phosphate across the inner membrane (33, 34) induced hexose phosphate re-uptake via the uhpBC two-component system and *uhpT* hexose phosphate transporter (30–32). A) A network map and regulatory schematic of the reactions into and out of the g6p node. The reaction in red is removed through the PGI KO. B) A mechanistic schematic of the uhpBC two-component system that sensed periplasmic hexose phosphate. The transcription factor UphA positively upregulated the expression of the hexose phosphate importer uhpT. C) Metabolite, expression, and flux levels near the node of perturbation. Abnormal elevations in glucose 6-phosphate (g6p) and imbalance of the glycolytic intermediates in pgi were found to induce a sugar phosphate toxicity response sensed through sgrR and mediated through the action of the small RNA sgrS (6–8) (Panels A-B). D-E) Regulatory schematic of sgrR and sgrST-setA operons. Regulatory schematic of genes subjected to transcriptional activation or attenuation by the small RNA sgrS (7, 20, 55, 56). F) Gene expression profiles of sugar phosphate response genes. Note the elevations in g6p and corresponding upregulation of *sgrS* in response to activation of SgrR by g6p that is consistent with the literature (7, 20, 55, 56). Metabolite concentrations are derived from averages taken from triplicate cultures that were analyzed in duplicate (n=6). Gene expression values are derived from averages of biological duplicates.

### Imbalances in central carbon intermediates induced TRN responses that were consistent with the literature

Many *E. coli* TFs are activated by metabolites (22–26). Changes in metabolite levels were found to trigger myriad TRN responses as measured by changes in expression profiles associated and also consistent with well-known TF regulons. Many of these expression changes appeared to conflict with optimal fitness. One example of a counterproductive TF response was the activation of CRP by elevated cAMP levels. Such CRP activation generated a cascade of responses in alternate carbon metabolism operons(27, 28). These operons pertained to import and catabolism of sugars such as glycerol, maltose, mannose, etc., that were not present in the medium (Supplementary data). Specifically, the *glp* regulon required for glycerol import and catabolism was upregulated by CRP-cAMP(29). This hard-wired regulation led to massive up-regulation of the *glp* regulon in uPgi which could potentially have lead to counterproductive allocation of the proteome to glycerol metabolism.

Interestingly, the hexose phosphate importer, *uhpT*, was overexpressed in uPgi compared to Ref. High periplasmic g6p plausibly activated the uhpAB two-component system, which in turn up-regulates expression of the hexose phosphate importer *uhpT(30–32)*. This result suggests that the concentration build-up in uPgi was so great that g6p spilled over into the periplasmic space(33, 34) (Fig. 3, upper panels). Increased expression of *uhpT* could have generated a loop whereby excessive g6p that spilled into the periplasmic space would be re-imported into the cytosol. The transcriptional attenuation of *ptsG* by *sgrS* may act to compensate for this futile cycle.

### Imbalances in central carbon intermediates was mirrored in amino acid pools

In addition to regulatory shifts, biomass components directly reflected the levels of their biosynthetic precursors (Fig. 3B-C). The aromatic amino acids L-tyrosine (tyr-L), L-phenylalanine (phe-L), and L-tryptophan (trp-L) are derived from phosphoenolpyruvate (pep). A decrease in pep levels in uPgi and and a rise in pep levels in ePgi strains was mirrored by all three of the amino acids (Fig. 3C). Conversely, an increase in ribose 5 phosphate (r5p) levels in uPgi and return of r5p levels in ePgi strains was mirrored by the downstream amino acid L-histidine (his-L) and nucleotides IMP and UMP (Fig 3B).

Similar trends were found for amino acid and precursor pairs L-serine (ser-L) and 2-phosphogluconate (2pg), L-aspartate (asp-L) and oxaloacetate (oaa), L-alanine (ala-L) and pyruvate (pyr), and L-glutamate (glu-L), L-glutamine (glu-L), and alpha-ketoglutarate (akg), respectively (Table S3). Perturbations in the distribution and abundance of proteogenic amino acids have been shown to alter protein synthesis rates leading to a drop in the growth rate(35–39). The drop in the growth rate of the uPgi strain and regain of fitness in the ePgi strains provide evidence that an imbalance in glycolytic intermediates directly alter growth rate via manipulating proteogenic amino acid levels.

The regulatory response to elevated g6p levels and the relationship between biomass components and their precursors reflected the importance of balancing glycolytic, PPP, and TCA cycle intermediates to maintain balanced ratios of amino acids levels for protein biosynthesis and purines/pyrimidines for nucleotide biosynthesis.

### Mutations that targeted alternative glucose import systems corrected TRN responses, and helped to rebalance glycolytic intermediate levels

Major alterations in expression profiles were found in the evolved KO strains that correlated with mutations in TFs. These included mutations to *galR* (Fig. 5) and *malT* (Fig. 6) in the ePgi strains. A 22 nucleotide deletion in the small molecule binding domain of *galR* in ePgi07 appears to negate repression of *galR* controlled operons (Fig. 5). These include *galETKM, galP*, and *mglBAC* that encode enzymes for galactose catabolism, symport, and ABC transport, respectively(40). These operons are also regulated by CRP-cAMP, and were not expressed in Ref. The galactose importers have lesser affinity for the transport of glucose, which may give ePgi07 an additional route to import and catabolize glucose from the environment. In addition, the mutation may have aided in conserving pep for aromatic amino acid production, which was limiting fitness in all of the pgi strains (as discussed previously).

**Fig. 5.**
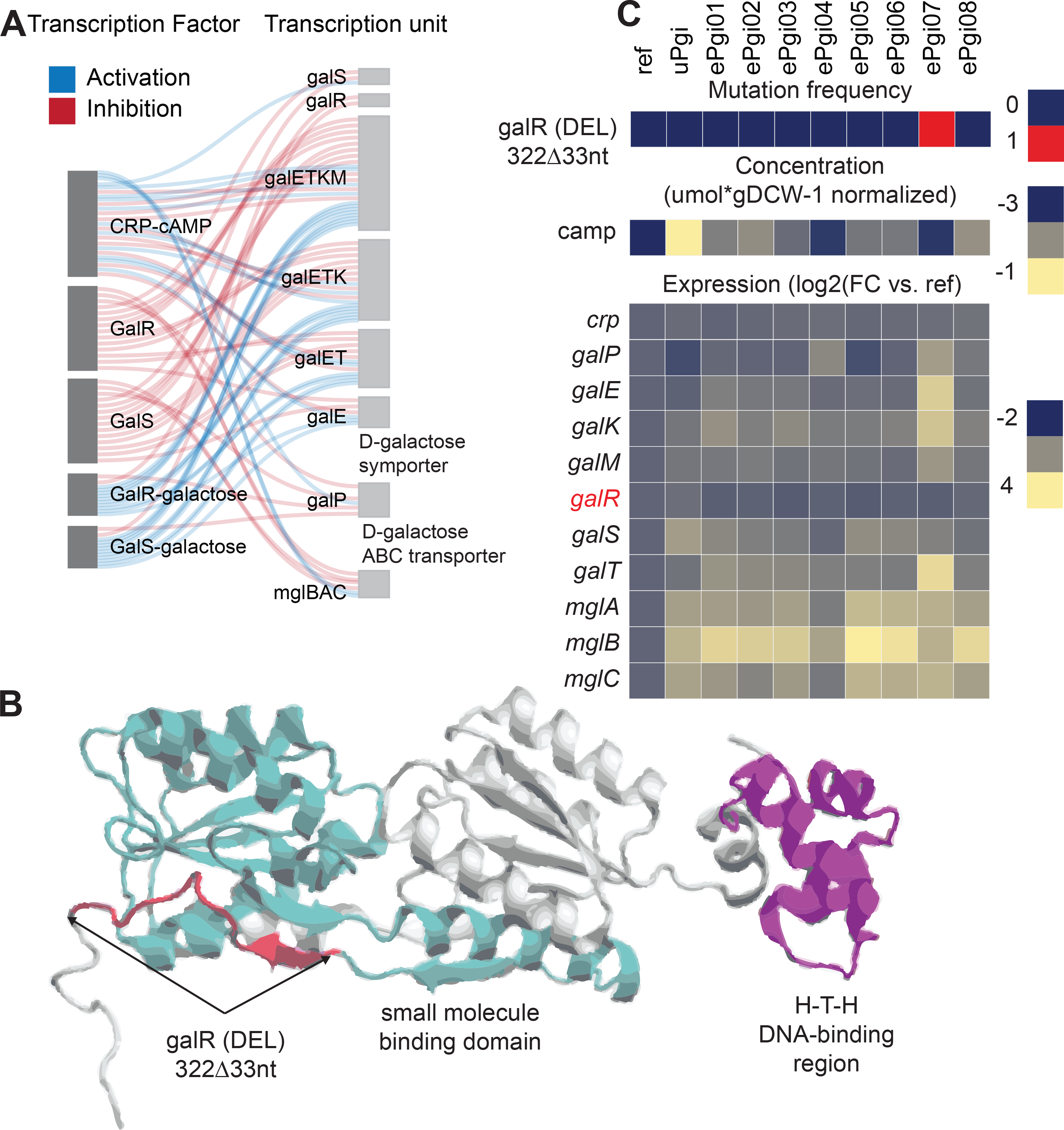
An inframe 33 nucleotide deletion (DEL) that removed 11 amino acids in the small molecule binding domain of galR negates galR repression in ePgi07 (Panels A-C). A) Regulatory network specifically controlled by cAMP-CRP, galR, and galS (57–59). cAMP-CRP can both positively and negatively regulate the expression of *galR, galS, galETKM, galP*, and *mglBAC;* GalR and GalS act as repressors; and GalR and GalS bound to galactose active primarily as activators. B) Crystal structure of the galR transcription factor (60). The position of the deletion is highlighted in red, the small molecule binding domain is highlighted in cyan, and the H-T-H DNA-binding region is highlighted in magenta. C) Mutation frequency for galR, metabolite concentration for cAMP, and expression profiles of galR controlled operons. Note the increased expression of *galP* and *galETKM* in ePgi07. Metabolite concentrations are derived from averages taken from triplicate cultures that were analyzed in duplicate (n=6). Gene expression values are derived from averages of biological duplicates.

**Fig. 6.**
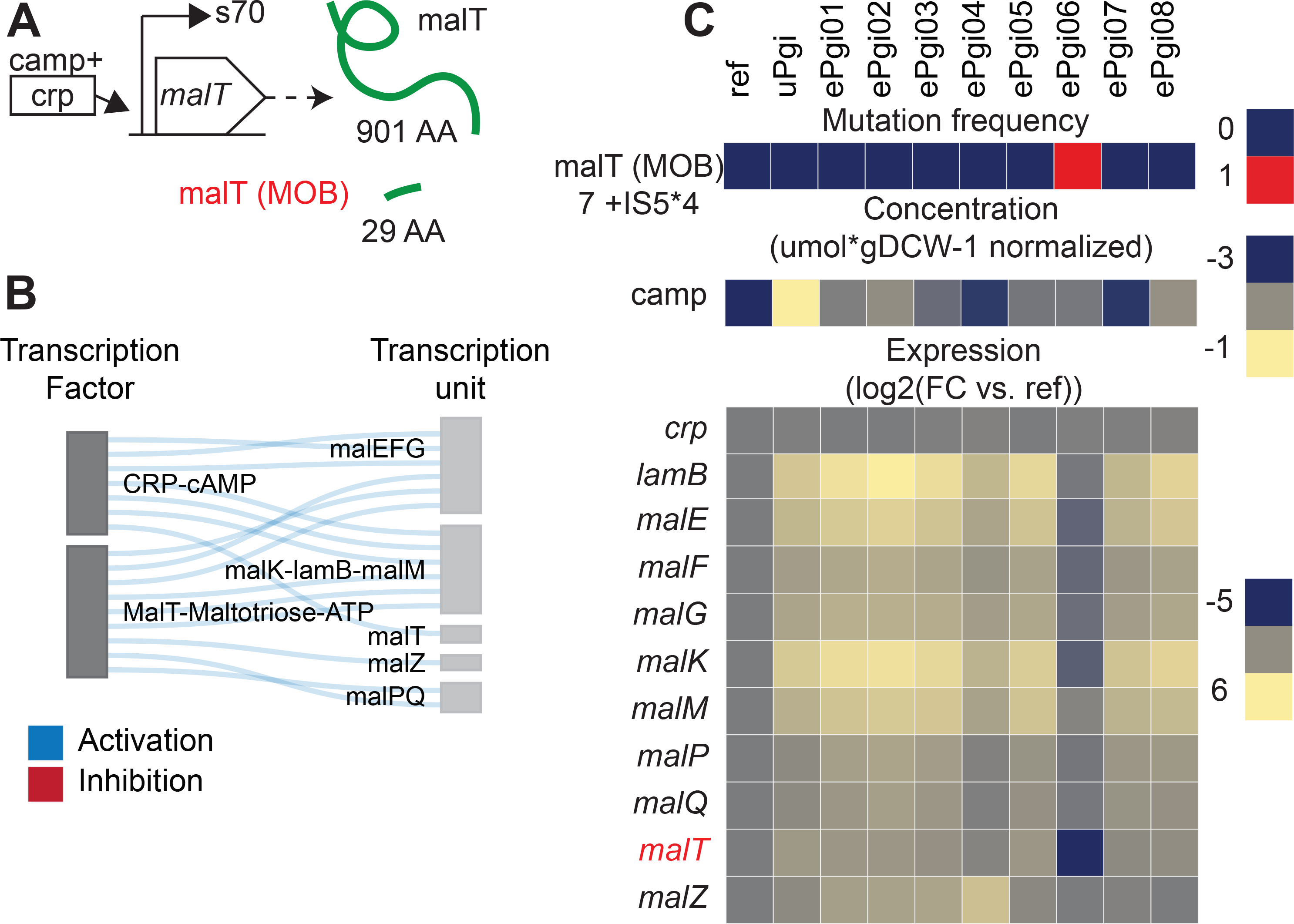
A mobile element insertion (MOB) that truncated the MalT TF in ePgi06 was found that appeared to silence expression of MalT controlled operons. A) Schematic of the malT operon(41) and truncated malT peptide. The mobile element insertion introduced a stop codon that reduced the MalT peptide from 901 amino acids to 29 amino acids. All binding-domains and catalytic sites were cleaved (42, 43). B) Operons controlled by malT(41). All regulators except malT and CRP-cAMP have been omitted. MalT controlled genes are involved in glycogen turnover, and may give ePgi06 an advantage in controlling the levels of hexose phosphates that are converted to and broken down from glycogen. C) Mutation frequency of malT, metabolite concentration for cAMP, and expression profiles for malT and malT regulated genes. Note the significantly repressed gene expression levels of MalT controlled genes in ePgi06. Metabolite concentrations are derived from averages taken from triplicate cultures that were analyzed in duplicate (n=6). Gene expression values are derived from averages of biological duplicates.

A mobile element insertion (MOB) that truncated the MalT TF in ePgi06 was found that appeared to silence expression of MalT(41) controlled operons (Fig. 6). The MOB introduced a stop codon that reduced the MalT peptide from 901 amino acids to 29 amino acids. All binding-domains and catalytic sites were cleaved(42, 43). MalT controlled genes are involved in glycogen turnover, and may give ePgi06 an advantage in controlling the levels of hexose phosphates that are converted to and broken down from glycogen.

### An imbalance in redox carriers was compensated for by shifts in hydrogenase flux and buffered by glutathione

Genome-scale MFA(18) confirmed that removal of the *pgi* gene diverted all upper glycolytic flux into the oxidative pentose phosphate pathway (oxPPP) (Fig. 2). The loss of PGI resulted in a 556.7% increase in flux per mol of glucose towards the oxPPP in uPgi (14.4 and 94.7% flux per mol of glucose in ref and uPgi, respectively). Note that as discussed previously, a large portion of the flux into the oxPPP was diverted down the ED pathway after the first NADPH generating step to avoid generating additional NADPH via the second NADPH generating step of the oxPPP (Fig. 2). Rearrangement of flux through hydrogenases to compensate for the increased flux towards NADPH generation were found (Fig. 2). Notable is the reversed utilization of the transhydrogenases from net NADPH to NADH generation in uPgi and ePgi strains (approximately -8 fold change in uPgi and ePgi strains in NADPH generation through THD2pp, and -3 fold change in uPgi and -1 to 4 fold change in ePgi strains in NADH generation through NADTRHD). Other hydrogenases significantly altered include serine dehydrogense (LSERDHr, note the altered levels of ser-L mentioned previously), as well as isocitrate dehydrogenase (ICD) and glutamate synthase (GLUSy, note the altered levels of akg, gln-L, and glu-L mentioned previously).

The increased flux through the oxPPP would generate an increased abundance of NADPH and thus a redox imbalance. LC-MS/MS(44) was used to quantify the absolute metabolite concentrations of the redox carriers. While major shifts in the redox carriers were found (Fig. 3A), a statistically significant change in NADPH between Ref and uPgi was not found. However, statistically significant changes in NADP and reduced and oxidized Glutathione were found. This indicates a potential rapid buffering of NADPH by the Glutathione via glutathione reductase (GTHOr) (Fig. 3A).

### High NADPH promoted activation of oxidative stress responders

Mutations in *soxR* in ePgi02 and *rseC* in ePgi01 were found that altered the expression of oxidative stress genes (Fig. 7). The *soxR* mutation cleaved the Fe-S cluster binding site of the SoxR peptide (Fig. 7C). Cleavage of the Fe-S cluster does not affect DNA-binding, but transcriptional activation of target genes *soxS* and *fumC* and transcriptional deactivation of *soxR* are impaired(45–47). *soxR* was up-regulated in ePgi02, which indicated that the mutation negated *soxR’s* self-regulation.

**Fig. 7.**
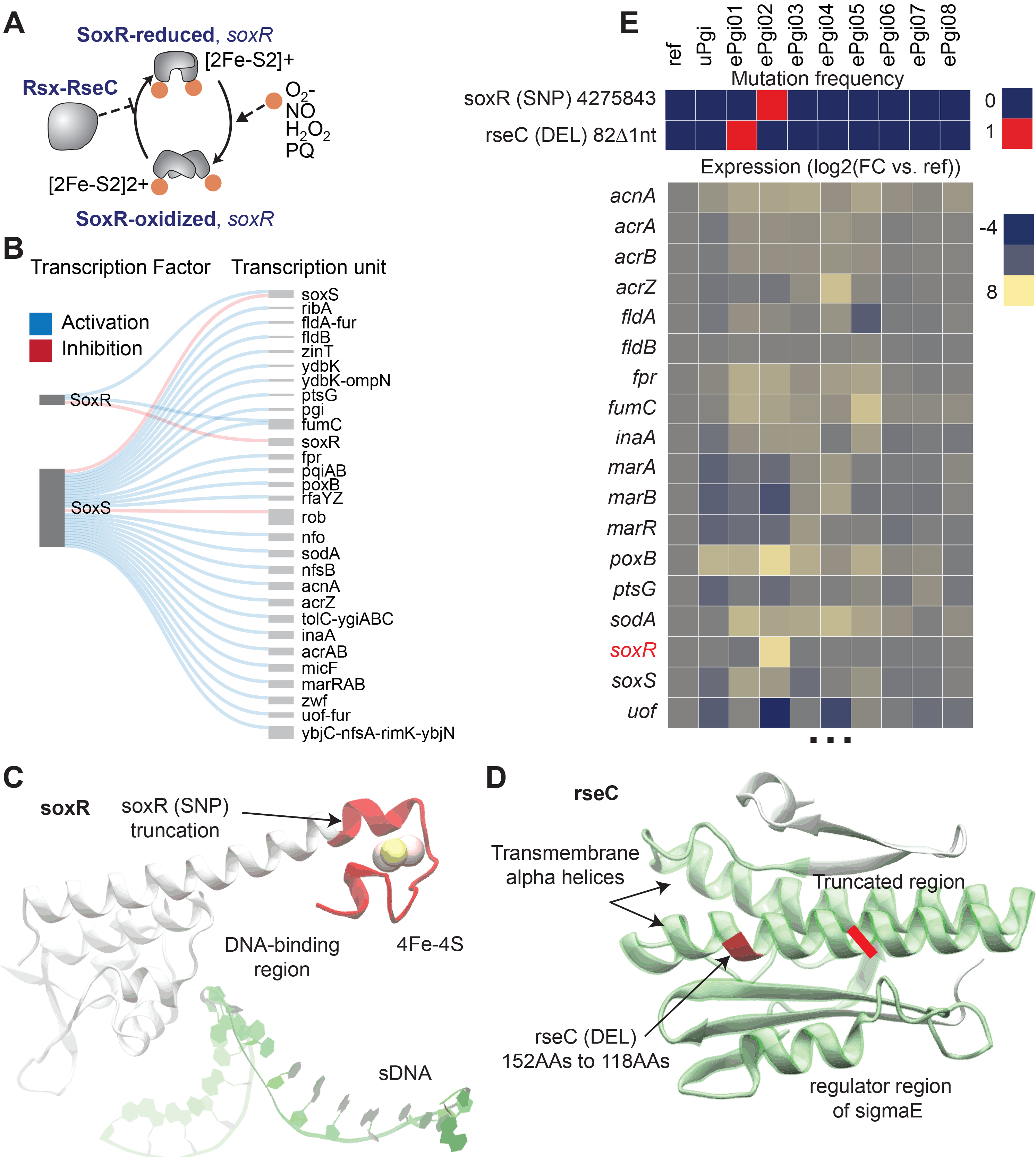
Mutations in *soxR* and *rseC* that altered the expression of oxidative stress genes. A) protein-protein interaction schema between SoxR and Rsx-RseC. In the reduced form, the iron sulfur clusters of the SoxR homodimers sense the presence of free radicals and ROS(61, 62). The oxidation of the iron sulfur clusters by free radicals and ROS induces a conformation changes from the inactive form to the active form(63). While both reduced and inactive form and oxidized and active forms of SoxR are capable of binding DNA, only the active form is capable of activating or inhibiting transcription(45, 64–67). The Rsx-RseC complex prevents reduction and inactivation of SoxR(48). B) Regulatory schematic of a subset of SoxR and SoxS controlled operons. C) Crystal structure of SoxR (68). The *soxR* SNP cleaved the Fe-S cluster binding site of the SoxR peptide. The SoxR DNA-binding region in proximity to single stranded DNA (sDNA) is shown below. D) Crystal structure of rseC. The *rseC* mutation cleaved a large portion of the transmembrane helix region that may affect Rsx-RseC complex formation or activity. The mutated and/or cleaved residues are shown in red. E) Mutation frequency and gene expression profiles. Gene expression values are derived from averages of biological duplicates.

The *rseC* mutation cleaved a large portion of the transmembrane helix region (Fig. 7D) that may affect Rsx-RseC complex formation or activity. *rseC* mutants were found previously to exhibit constitutive *soxS* expression by preventing the Rsx-RseC complex to inhibit reduction and inactivation of SoxR(48) (Fig. 7A). *soxS*, as well as many of its downstream activation targets including *acrA, acrB, fldA, fpr, inaA*, and *sodA*, were up-regulated in ePgi01, which indicated that the mutation promotes expression of soxS. The Rsx-RseC complex prevents reduction and inactivation of SoxR(48).

### Mutations in transhydrogenases helped to alleviate redox imbalance

Mutations selected during adaptive evolution also introduced innovations that targeted metabolic network elements involved in NADPH production. Mutated transhydrogenases included *sthA, pntB, icd*, and *zwf*. The soluble and membrane bound transhydrogenases act to interconvert NADP(H) and NAD(H) (49, 50) (Fig. 8). Mutations in the soluble *sthA (49)* and membrane bound *pntB (50)* transhydrogenases in ePgi04 and ePgi07, respectively, were found (Fig. 8). The *sthA* mutation appeared near the dimerization domain, and may affect enzyme complex formation. The *pntB* mutation appeared in the transmembrane region, and may affect catalytic activity or membrane association. It has been demonstrated that altered activity of *sthA* and *pntAB* confers a fitness advantage in *pgi* mutant strains by rebalancing the ratios of NADH to NADPH (49, 50). Interestingly, mutations in *sthA* and *pntB* were selected for in previous evolutions of a *pgi* strain (3). This observation provides further evidence that the *sthA* and *pntB* mutations provided a fitness advantage to ePgi04 and ePgi07 by rebalancing the ratios of NADH to NADPH via modulating the activity of the transhydrogenases. Note that ePgi04 and ePgi07 were also found to have the highest increase in flux through the soluble transhydrogenases.

**Fig. 8.**
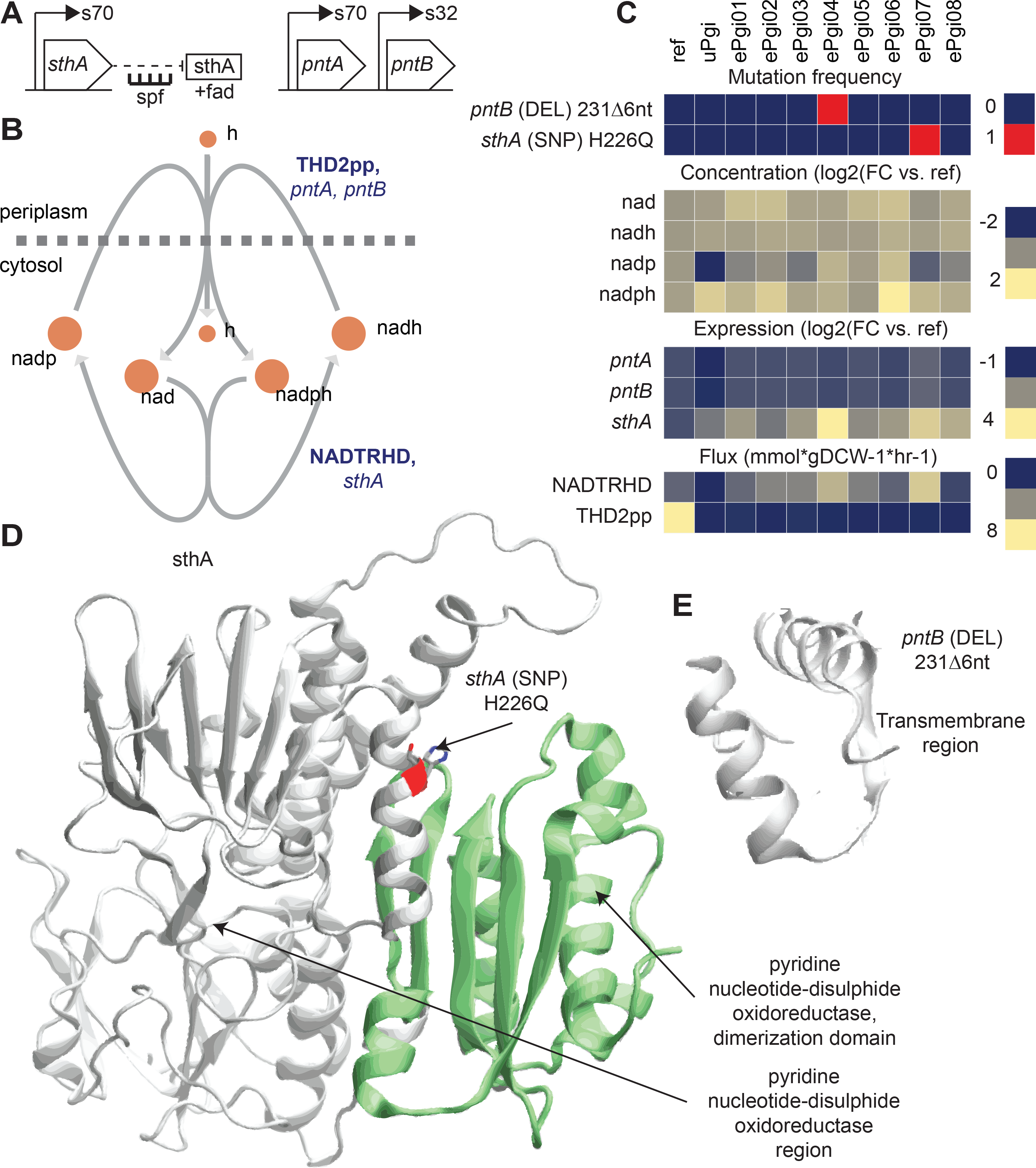
Mutations in the soluble *sthA (49)* and membrane bound *pntB (50)* transhydrogenases that potentially aid in balancing NAD(P)(H) cofactors. A) Schematic of the *sthA* and *pntAB* operons. B) Network diagrams of the soluble pyridine nucleotide transhydrogenase (NADTRHD) reaction catalyzed by *sthA* and the membrane bound pyridine nucleotide transhydrogenase (THD2pp) reaction catalyzed by *pntAB*. C) Mutation frequency and metabolite and expression levels near the genes. D) The *sthA* mutation in ePgi04 appeared near the dimerization domain, and may affect enzyme complex formation. E) The *pntB* mutation in ePgi07 appeared in the transmembrane region, and may affect catalytic activity or membrane association. It has been demonstrated that altered activity of *sthA* and *pntAB* confers a fitness advantage in pgi mutant strains by rebalancing the ratios of NADH to NADPH (49, 50). Metabolite concentrations are derived from averages taken from triplicate cultures that were analyzed in duplicate (n=6). Gene expression values are derived from averages of biological duplicates.

Isocitrate dehydrogenase (ICD) catalyzes the conversion of isocitrate (icit) to 2-oxoglutarate (akg) while reducing NADP+ to NADPH (Fig. 9). Activity of ICD also regulates the flux split between the full TCA cycle and the glyoxylate shunt(51–53). A point mutation at the 395 residue that changed the amino acid from positively charged (L-arginine) to negatively charged (L-cysteine) in ICD was found in all ePgi replicates except replicate 7 (Fig 9). The mutation occurs 4 Angstroms from the phosphate moiety of NADP. The 395 residue has been shown to be directly involved in NADP-binding (54), and appears to allow the mutated enzyme to utilize NAD as a cofactor. The mutation was found to redirect flux through the glyoxylate shunt instead of the TCA cycle, and may provide a fitness advantage to the ePgi strains by limiting the production of NADPH in the TCA cycle.

**Fig. 9.**
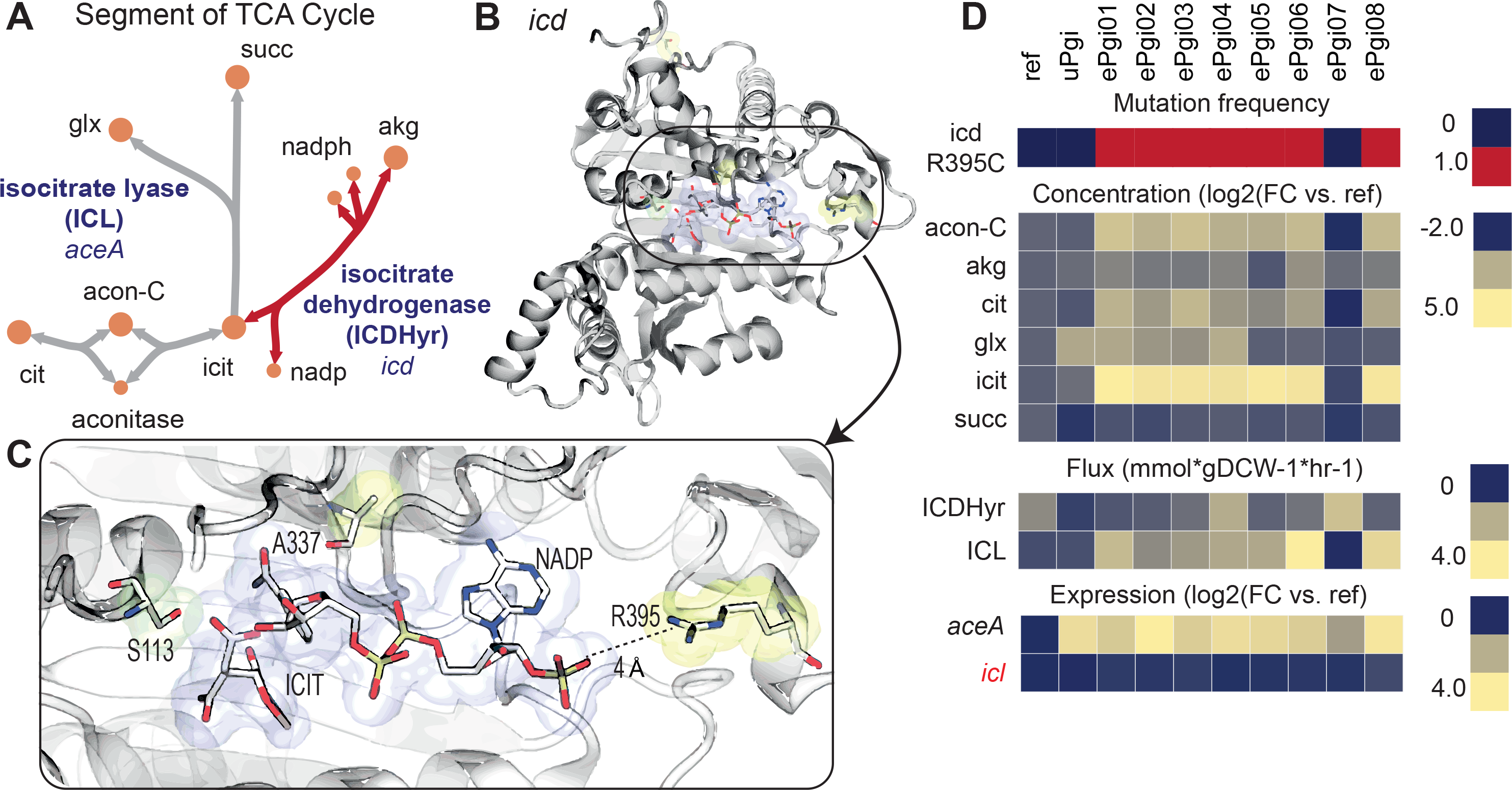
A beneficial mutation that rewired the TCA cycle via a cofactor usage swap in isocitrate dehydrogenase (ICD) aided in alleviating the excessive conversion of NADP to NADPH. A) Network schematic of a segment of the TCA cycle. The reaction in red is catalyzed by ICD where mutations fix during ALE. B) Crystal structure of ICD. The mutated amino acids are highlighted in yellow. C) Zoom in on the active site of isocitrate dehydrogenase showing the proximity of the mutated amino acid to the phosphate group of NADP. The mutation occurs 4 Angstroms from the phosphate moiety of NADPH. The 395 residue has been shown to be directly involved in NADPH-binding (54), and appears to allow the mutated enzyme to utilize NADH as a cofactor. D) Mutation frequency and metabolite, expression, and flux levels near the mutated gene. System components near the ICDHyr reaction in the ICD mutant strains are significantly changed. Metabolite concentrations are derived from averages taken from triplicate cultures that were analyzed in duplicate (n=6). Flux levels are derived from averages taken from triplicate cultures that were analyzed in duplicate (n=6). Gene expression values are derived from averages of biological duplicates.

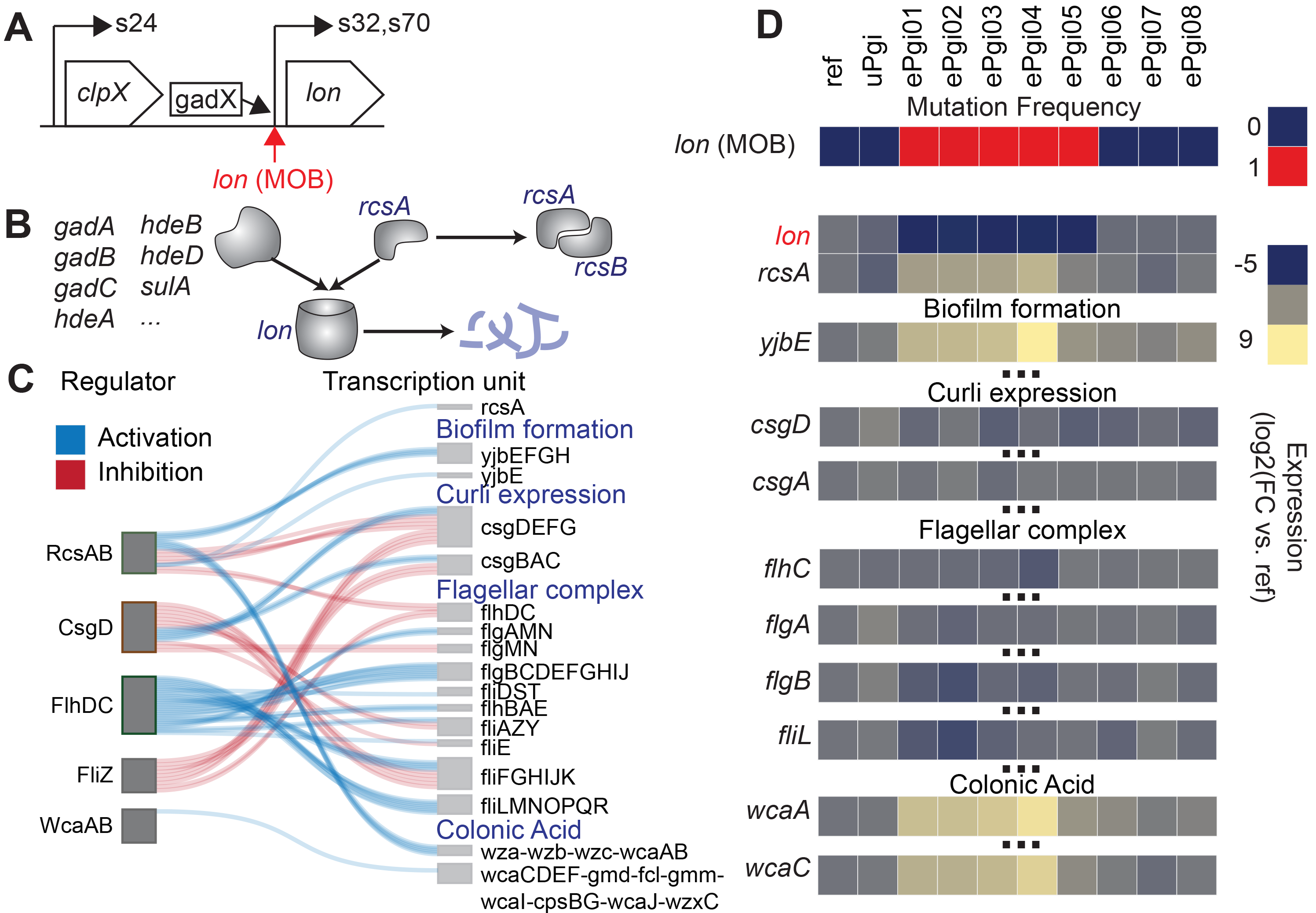

### Metabolome, fluxome, transcriptome, and genome were unique in each ePgi strain

While the ePgi strains were not able to recover the fitness of Ref, they were able to recover the fitness of wild-type MG1655. Many intermediate and cofactor levels, including g6p, remained perturbed in ePgi strains to varying degrees. However, the majority of initially elevated transcription involving sugar phosphate stress, carbon catabolite repression, the *uhpT* transporter, and other general stress responders in uPgi were dampened or completely shut down in ePgi strains. This indicated that the TRN had evolved to cope with the changed metabolome. Many of these changes to the TRN could be directly attributed to mutations.

The recovery to wild-type levels was in part made possible by a complete rewiring of central carbohydrate metabolic flux splits (Fig. 3). The rewiring differed substantially between ePgi strains. While an increase in flux through GND and a decrease in flux through the ED pathway occurred in all ePgi endpoints, the flux through each pathway differed substantially. GND flux varied from 389.9 to 604.6% increase per mol glucose compared to Ref, and ED pathway flux varied from 150.9 to 4463.8% increase per mol glucose compared to Ref. NonOxPPP flux varied substantially between the strains, even altering between net forward and reverse utilization of TALA. Of particular note, flux through PPC was regained in ePgi04 and ePgi07, and flux through the glyoxylate shunt was lost in ePgi07. The later differences in particular correlate with mutations in the transhydrogenases and *icd*, respectively.

## Conclusion

Loss of *pgi* induced massive perturbations to the metabolome, fluxome, and transcriptome that led to a greatly retarded growth rate. In contrast to previous work (5), it was found that the loss of *pgi* induced major changes in all -omics data measured not just local to the perturbation, but in distal network locations. Flux rerouting to compensate for the loss of *pgi* imbalanced PPP and glycolytic intermediate levels, which lead to a sugar phosphate stress response. The deleterious effects of this response were attributed to a misallocation of protein, a deleterious cycle of re-import of hexose phosphate, and alterations in the distributions and amount of proteogenic amino acids and nucleotides. Redistribution of glycolytic flux into the oxPPP generated an overabundance of NADPH, which led to a redox imbalance.

ALE selected for mutations that helped to alleviate redox and glycolytic intermediate imbalances. Differences in the metabolome and mutation landscape led to a diversity of expression and fluxome profiles in ePgi strains. The multitude of hydrogenases and routes to generate glycolytic intermediates allowed for regulatory and metabolic flexibility in overcoming redox and glycolytic intermediate imbalance. The diversity in mutation fixed and the concomitant emergence of multiple optimal phenotypes was a manifestation of this flexibility.

## Contributions

D.M. designed the experiments; generated the strains; conducted all aspects of the metabolomics, fluxomics, phenomics, transcriptomics, and genomics experiments; performed all multi-omics statistical, graph, and modeling analyses; and wrote the manuscript. T.E.S. ran the ALE experiments. E.B. assisted with structural analysis. R.S. processed the DNA and RNA samples. S.X. assisted with metabolomics and fluxomics data collection, sample processing, and peak integration. Y.H. assisted with fluxomics data collection and sample processing. A.M.F designed and supervised the evolution experiments, and contributed to the data analysis and the manuscript. B.O.P conceived and outlined the study, supervised the data analysis, and co-wrote the manuscript.

## Acknowledgements

We thank José Utrilla for helpful discussion and guidance when implementing the knockouts in the pre-evolved strain. We thank Jamey Young for helpful discussions throughout the MFA analysis. We thank Laurence Yang for helpful discussions regarding optimization and statistical analysis. This work was funded by the Novo Nordisk Foundation Grant Number NNF10CC1016517.

## Competing financial interests

The authors declare no competing financial interests.

## References

1. Usui Y, Hirasawa T, Furusawa C, Shirai T, Yamamoto N, Mori H, Shimizu H. 2012. Investigating the effects of perturbations to pgi and eno gene expression on central carbon metabolism in Escherichia coli using (13)C metabolic flux analysis. Microb Cell Fact 11:87.

2. Ahn J, Chung BKS, Lee D-Y, Park M, Karimi IA, Jung J-K, Lee H. 2011. NADPH-dependent pgi-gene knockout Escherichia coli metabolism producing shikimate on different carbon sources. FEMS Microbiol Lett 324:10–16.

3. Charusanti P, Conrad TM, Knight EM, Venkataraman K, Fong NL, Xie B, Gao Y, Palsson BØ. 2010. Genetic basis of growth adaptation of Escherichia coli after deletion of pgi, a major metabolic gene. PLoS Genet 6:e1001186.

4. Fong SS, Nanchen A, Palsson BO, Sauer U. 2006. Latent pathway activation and increased pathway capacity enable Escherichia coli adaptation to loss of key metabolic enzymes. J Biol Chem 281:8024–8033.

5. Ishii N, Nakahigashi K, Baba T, Robert M, Soga T, Kanai A, Hirasawa T, Naba M, Hirai K, Hoque A, Ho PY, Kakazu Y, Sugawara K, Igarashi S, Harada S, Masuda T, Sugiyama N, Togashi T, Hasegawa M, Takai Y, Yugi K, Arakawa K, Iwata N, Toya Y, Nakayama Y, Nishioka T, Shimizu K, Mori H, Tomita M. 2007. Multiple high-throughput analyses monitor the response of E. coli to perturbations. Science 316:593–597.

6. Richards GR, Patel MV, Lloyd CR, Vanderpool CK. 2013. Depletion of glycolytic intermediates plays a key role in glucose-phosphate stress in Escherichia coli. J Bacteriol 195:4816–4825.

7. Vanderpool CK, Gottesman S. 2004. Involvement of a novel transcriptional activator and small RNA in post-transcriptional regulation of the glucose phosphoenolpyruvate phosphotransferase system. Mol Microbiol 54:1076–1089.

8. Vanderpool CK, Gottesman S. 2007. The novel transcription factor SgrR coordinates the response to glucose-phosphate stress. J Bacteriol 189:2238–2248.

9. Holmgren A. 1976. Hydrogen donor system for Escherichia coli ribonucleoside-diphosphate reductase dependent upon glutathione. Proceedings of the National Academy of Sciences 73:2275–2279.

10. Lin Z, Xu Z, Li Y, Wang Z, Chen T, Zhao X. 2014. Metabolic engineering of Escherichia coli for the production of riboflavin. Microb Cell Fact 13:104.

11. Tenaillon O, Barrick JE, Ribeck N, Deatherage DE, Blanchard JL, Dasgupta A, Wu GC, Wielgoss S, Cruveiller S, Médigue C, Schneider D, Lenski RE. 2016. Tempo and mode of genome evolution in a 50,000-generation experiment. Nature 536:165–170.

12. Plucain J, Hindré T, Le Gac M, Tenaillon O, Cruveiller S, Médigue C, Leiby N, Harcombe WR, Marx CJ, Lenski RE, Schneider D. 2014. Epistasis and Allele Specificity in the Emergence of a Stable Polymorphism in Escherichia coli. Science 343:1366–1369.

13. Dragosits M, Mattanovich D. 2013. Adaptive laboratory evolution -- principles and applications for biotechnology. Microb Cell Fact 12:64.

14. Chou H-H, Marx CJ, Sauer U. 2015. Transhydrogenase promotes the robustness and evolvability of E. coli deficient in NADPH production. PLoS Genet 11:e1005007.

15. Nakahigashi K, Toya Y, Ishii N, Soga T, Hasegawa M, Watanabe H, Takai Y, Honma M, Mori H, Tomita M. 2009. Systematic phenome analysis of Escherichia coli multiple-knockout mutants reveals hidden reactions in central carbon metabolism. Mol Syst Biol 5:306.

16. LaCroix RA, Sandberg TE, O’Brien EJ, Utrilla J, Ebrahim A, Guzman GI, Szubin R, Palsson BO, Feist AM. 2015. Use of Adaptive Laboratory Evolution To Discover Key Mutations Enabling Rapid Growth of Escherichia coli K-12 MG1655 on Glucose Minimal Medium. Appl Environ Microbiol 81:17–30.

17. Sandberg TE, Pedersen M, LaCroix RA, Ebrahim A, Bonde M, Herrgard MJ, Palsson BO, Sommer M, Feist AM. 2014. Evolution of Escherichia coli to 42 °C and subsequent genetic engineering reveals adaptive mechanisms and novel mutations. Mol Biol Evol 31:2647–2662.

18. McCloskey D, Young JD, Xu S, Palsson BO, Feist AM. 2016. Modeling Method for Increased Precision and Scope of Directly Measurable Fluxes at a Genome-Scale. Anal Chem 88:3844–3852.

19. Bak G, Lee J, Suk S, Kim D, Young Lee J, Kim K-S, Choi B-S, Lee Y. 2015. Identification of novel sRNAs involved in biofilm formation, motility, and fimbriae formation in Escherichia coli. Sci Rep 5:15287.

20. Kimata K, Tanaka Y, Inada T, Aiba H. 2001. Expression of the glucose transporter gene, ptsG, is regulated at the mRNA degradation step in response to glycolytic flux in Escherichia coli. EMBO J 20:3587–3595.

21. Vanderpool CK. 2007. Physiological consequences of small RNA-mediated regulation of glucose-phosphate stress. Curr Opin Microbiol 10:146–151.

22. Gama-Castro S, Salgado H, Santos-Zavaleta A, Ledezma-Tejeida D, Muñiz-Rascado L, García-Sotelo JS, Alquicira-Hernández K, Martínez-Flores I, Pannier L, Castro-Mondragón JA, Medina-Rivera A, Solano-Lira H, Bonavides-Martínez C, Pérez-Rueda E, Alquicira-Hernández S, Porrón-Sotelo L, López-Fuentes A, Hernández-Koutoucheva A, Del Moral-Chávez V, Rinaldi F, Collado-Vides J. 2016. RegulonDB version 9.0: high-level integration of gene regulation, coexpression, motif clustering and beyond. Nucleic Acids Res 44:D133–43.

23. Cho S, Cho Y-B, Kang TJ, Kim SC, Palsson B, Cho B-K. 2015. The architecture of ArgR-DNA complexes at the genome-scale in Escherichia coli. Nucleic Acids Res 43:3079–3088.

24. Federowicz S, Kim D, Ebrahim A, Lerman J, Nagarajan H, Cho B-K, Zengler K, Palsson B. 2014. Determining the control circuitry of redox metabolism at the genome-scale. PLoS Genet 10:e1004264.

25. Kim D, Hong JS-J, Qiu Y, Nagarajan H, Seo J-H, Cho B-K, Tsai S-F, Palsson BØ. 2012. Comparative analysis of regulatory elements between Escherichia coli and Klebsiella pneumoniae by genome-wide transcription start site profiling. PLoS Genet 8:e1002867.

26. Cho B-K, Federowicz S, Park Y-S, Zengler K, Palsson BØ. 2011. Deciphering the transcriptional regulatory logic of amino acid metabolism. Nat Chem Biol 8:65–71.

27. You C, Okano H, Hui S, Zhang Z, Kim M, Gunderson CW, Wang Y-P, Lenz P, Yan D, Hwa T. 2013. Coordination of bacterial proteome with metabolism by cyclic AMP signalling. Nature 500:301–306.

28. Hermsen R, Okano H, You C, Werner N, Hwa T. 2015. A growth◻rate composition formula for the growth of E. coli on co◻utilized carbon substrates. Mol Syst Biol 11:801.

29. Larson TJ, Cantwell JS, van Loo-Bhattacharya AT. 1992. Interaction at a distance between multiple operators controls the adjacent, divergently transcribed glpTQ-glpACB operons of Escherichia coli K-12. J Biol Chem 267:6114–6121.

30. Weston LA, Kadner RJ. 1988. Role of uhp genes in expression of the Escherichia coli sugar-phosphate transport system. J Bacteriol 170:3375–3383.

31. Dahl JL, Wei BY, Kadner RJ. 1997. Protein phosphorylation affects binding of the Escherichia coli transcription activator UhpA to the uhpT promoter. J Biol Chem 272:1910–1919.

32. Maloney PC, Ambudkar SV, Anatharam V, Sonna LA, Varadhachary A. 1990. Anion-exchange mechanisms in bacteria. Microbiol Rev 54:1–17.

33. Bolten CJ, Kiefer P, Letisse F, Portais J-C, Wittmann C. 2007. Sampling for metabolome analysis of microorganisms. Anal Chem 79:3843–3849.

34. Link H, Anselment B, Weuster-Botz D. 2008. Leakage of adenylates during cold methanol/glycerol quenching of Escherichia coli. Metabolomics 4:240–247.

35. Stoebel DM, Dean AM, Dykhuizen DE. 2008. The cost of expression of Escherichia coli lac operon proteins is in the process, not in the products. Genetics 178:1653–1660.

36. Kaleta C, Schäuble S, Rinas U, Schuster S. 2013. Metabolic costs of amino acid and protein production in Escherichia coli. Biotechnol J 8:1105–1114.

37. Traxler MF, Chang D-E, Conway T. 2006. Guanosine 3’,5’-bispyrophosphate coordinates global gene expression during glucose-lactose diauxie in Escherichia coli. Proc Natl Acad Sci U S A 103:2374–2379.

38. Tao H, Bausch C, Richmond C, Blattner FR, Conway T. 1999. Functional genomics: expression analysis of Escherichia coli growing on minimal and rich media. J Bacteriol 181:6425–6440.

39. Akashi H, Gojobori T. 2002. Metabolic efficiency and amino acid composition in the proteomes of Escherichia coli and Bacillus subtilis. Proc Natl Acad Sci U S A 99:3695–3700.

40. Weickert MJ, Adhya S. 1993. The galactose regulon of Escherichia coli. Mol Microbiol 10:245–251.

41. Schlegel A, Böhm A, Lee S-J, Peist R, Decker K, Boos W. 2002. Network regulation of the Escherichia coli maltose system. J Mol Microbiol Biotechnol 4:301–307.

42. Liu P, Danot O, Richet E. 2013. A dual role for the inducer in signalling by MalT, a signal transduction ATPase with numerous domains (STAND). Mol Microbiol 90:1309–1323.

43. Danot O. 2010. The inducer maltotriose binds in the central cavity of the tetratricopeptide-like sensor domain of MalT, a bacterial STAND transcription factor. Mol Microbiol 77:628–641.

44. McCloskey D, Gangoiti JA, Palsson BO, Feist AM. 2015. A pH and solvent optimized reverse-phase ion-paring-LC-MS/MS method that leverages multiple scan-types for targeted absolute quantification of intracellular metabolites. Metabolomics 11:1338–1350.

45. Gaudu P, Weiss B. 1996. SoxR, a [2Fe-2S] transcription factor, is active only in its oxidized form. Proceedings of the National Academy of Sciences 93:10094–10098.

46. Bradley TM, Hidalgo E, Leautaud V, Ding H, Demple B. 1997. Cysteine-to-Alanine Replacements in the Escherichia Coli SoxR Protein and the Role of the [2Fe-2S] Centers in Transcriptional Activation. Nucleic Acids Res 25:1469–1475.

47. Tseng C-P, Yu C-C, Lin H-H, Chang C-Y, Kuo J-T. 2001. Oxygen- and Growth Rate-Dependent Regulation ofEscherichia coli Fumarase (FumA, FumB, and FumC) Activity. J Bacteriol 183:461–467.

48. Koo M-S, Lee J-H, Rah S-Y, Yeo W-S, Lee J-W, Lee K-L, Koh Y-S, Kang S-O, Roe J-H. 2003. A reducing system of the superoxide sensor SoxR in Escherichia coli. EMBO J 22:2614–2622.

49. Canonaco F, Hess TA, Heri S, Wang T, Szyperski T, Sauer U. 2001. Metabolic flux response to phosphoglucose isomerase knock-out in Escherichia coli and impact of overexpression of the soluble transhydrogenase UdhA. FEMS Microbiol Lett 204:247–252.

50. Sauer U, Canonaco F, Heri S, Perrenoud A, Fischer E. 2004. The soluble and membrane-bound transhydrogenases UdhA and PntAB have divergent functions in NADPH metabolism of Escherichia coli. J Biol Chem 279:6613–6619.

51. Walsh K, Koshland DE Jr. 1985. Branch point control by the phosphorylation state of isocitrate dehydrogenase. A quantitative examination of fluxes during a regulatory transition. J Biol Chem 260:8430–8437.

52. McKEE JS, Nimmo HG. 1989. Evidence for an arginine residue at the coenzyme-binding site of Escherichia coli isocitrate dehydrogenase. Biochem J 261:301–304.

53. LaPorte DC, Walsh K, Koshland DE Jr. 1984. The branch point effect. Ultrasensitivity and subsensitivity to metabolic control. J Biol Chem 259:14068–14075.

54. Zhu G, Golding GB, Dean AM. 2005. The selective cause of an ancient adaptation. Science 307:1279–1282.

55. Sun Y, Vanderpool CK. 2013. Physiological consequences of multiple-target regulation by the small RNA SgrS in Escherichia coli. J Bacteriol 195:4804–4815.

56. Bobrovskyy M, Vanderpool CK. 2016. Diverse mechanisms of post-transcriptional repression by the small RNA regulator of glucose-phosphate stress. Mol Microbiol 99:254–273.

57. Geanacopoulos M, Adhya S. 1997. Functional characterization of roles of GalR and GalS as regulators of the gal regulon. J Bacteriol 179:228–234.

58. Aki T, Adhya S. 1997. Repressor induced site-specific binding of HU for transcriptional regulation. EMBO J 16:3666–3674.

59. Semsey S, Krishna S, Sneppen K, Adhya S. 2007. Signal integration in the galactose network of Escherichia coli. Mol Microbiol 65:465–476.

60. Xu D, Zhang Y. 2013. Ab Initio structure prediction for Escherichia coli: towards genome-wide protein structure modeling and fold assignment. Sci Rep 3:1895.

61. Liochev SI, Fridovich I. 2011. Is superoxide able to induce SoxRS? Free Radic Biol Med 50:1813.

62. Lo F-C, Lee J-F, Liaw W-F, Hsu I-J, Tsai Y-F, Chan SI, Yu SS-F. 2012. The metal core structures in the recombinant Escherichia coli transcriptional factor SoxR. Chemistry 18:2565–2577.

63. Kobayashi K. 2017. Sensing Mechanisms in the Redox-Regulated, [2Fe-2S] Cluster-Containing, Bacterial Transcriptional Factor SoxR. Acc Chem Res 50:1672–1678.

64. Ding H, Demple B. 1996. Glutathione-mediated destabilization in vitro of [2Fe-2S] centers in the SoxR regulatory protein. Proceedings of the National Academy of Sciences 93:9449–9453.

65. Hidalgo E, Demple B. 1994. An iron-sulfur center essential for transcriptional activation by the redox-sensing SoxR protein. EMBO J 13:138–146.

66. Hidalgo E, Bollinger JM Jr, Bradley TM, Walsh CT, Demple B. 1995. Binuclear [2Fe-2S] clusters in the Escherichia coli SoxR protein and role of the metal centers in transcription. J Biol Chem 270:20908–20914.

67. Hidalgo E, Demple B. 1996. Activation of SoxR-dependent transcription in vitro by noncatalytic or NifS-mediated assembly of [2Fe-2S] clusters into apo-SoxR. J Biol Chem 271:7269–7272.

68. Watanabe S, Kita A, Kobayashi K, Miki K. 2008. Crystal structure of the [2Fe-2S] oxidative-stress sensor SoxR bound to DNA. Proc Natl Acad Sci U S A 105:4121–4126.

